# The Modulation of the Blood-Brain Barrier by Focused Ultrasound Stimulates Oligodendrogenesis

**DOI:** 10.1101/2025.05.21.655298

**Authors:** Kate Noseworthy, Joseph Silburt, Kullervo Hynynen, Isabelle Aubert

## Abstract

**Objective:** The current study aims to fill a gap in knowledge on the effects of focused ultrasound (FUS)-mediated blood-brain-barrier (BBB) modulation on the proliferation and development of oligodendrocyte progenitor cells (OPCs). Researchers established that FUS combined with intravenous microbubbles can modulate the BBB in a controlled, reversible, localized, and non-invasive manner to facilitate the delivery of intravenous therapeutics to the brain. Over a decade ago, we discovered that, even without intravenous therapeutics, FUS-BBB modulation stimulates elements of brain repair, including hippocampal neurogenesis.

**Methods:** In adult mice, FUS-BBB modulation was targeted unilaterally to the hippocampus and proliferation of OPCs was quantified at 1, 4, 7, and 10 days post-FUS. Mature oligodendrocytes were quantified at 30 days post-FUS. OPC proliferation was assessed at 7 days post-FUS, and mature oligodendrocytes at 30 days.

**Results:** The proliferation of hippocampal OPCs was increased by 6.8-fold and 2.3-fold between 1 and 4 days post-sonication, respectively, resulting in a 5.3-fold increase in mature oligodendrocytes one month later. To test the robustness of oligodendrogenesis following FUS-BBB modulation, the striatum was targeted as a second brain region with an independent experimental design. In line with hippocampal results, striatal FUS-BBB modulation promoted the generation of OPCs by 3.9-fold during the first week, leading to a 5.2-fold increase in oligodendrogenesis 30 days post-treatment.

**Interpretation:** We conclude that FUS-BBB modulation in the hippocampus and striatum promotes oligodendrogenesis by stimulating the proliferation of OPCs and being permissive to their maturation.

**Highlights:**

- Beyond the potential for the delivery of therapeutics to the brain, the modulation of the BBB by FUS can stimulate regenerative effects, including oligodendrogenesis.
- FUS-BBB modulation induced a significant proliferation of OPCs which resulted in increases in oligodendrogenesis of 5.3-fold in the hippocampus and 6.7-fold in the striatum

## 1. Introduction

The transient modulation of the blood-brain barrier (BBB) induced by the interactions of non-invasive focused ultrasound (FUS) with intravenous microbubbles [1] has been established as safe in animal models and in patients with glioblastoma [2–5], Alzheimer disease (AD) [6–9], amyotrophic lateral sclerosis [10] and Parkinson’s disease [11,12].

In addition to the delivery of intravenous therapeutics to the brain without surgical interventions [6,13–18], FUS-BBB modulation can trigger regenerative effects in the brain. For example, FUS-BBB modulation targeted to the hippocampus increases hippocampal neurogenesis through the proliferation of neural progenitor cells (NPCs), leading to the generation of immature neuroblasts and mature neurons [19–23]. FUS-BBB modulation has also been shown to prevent age-related dendrite loss [24] and to improve dendritic complexity and cognition in rodent models of AD [19]. The regenerative effects induced by FUS and microbubbles on different cell populations offers potential for multimodal treatment strategies for brain disorders.

The effects of FUS on oligodendrocytes and their progenitors, oligodendrocyte progenitor cells (OPCs), remain unexplored. OPCs are distributed throughout the central nervous system (CNS) [25–27]. Oligodendrogenesis, which refers to the process by which OPCs differentiate into mature oligodendrocytes, is critical for myelination in the physiological condition, and remyelination in pathological contexts. Oligodendrocytes not only contribute to myelin sheath formation but also participate in key myelin-independent aspects of development, function, and maintenance [28– 30].

This study aimed to test whether the transient modulation of the BBB by FUS and microbubbles in different brain regions promotes (1) the proliferation of OPCs in adult mice, and (2) the maturation and survival of newly-generated OPCs. The results established that FUS-BBB modulation in both the hippocampus and striatum enhanced the proliferation of OPCs and oligodendrogenesis. These findings highlight the potential of FUS-BBB modulation in creating a regenerative environment for brain repair, which could be of significance for brain disorders, including those with white matter pathologies such as AD [28,31] and multiple sclerosis [32].

### 2. Methods

#### 2.1. Animals

Male C57BL/6J mice (n=48) were used at 2.5 – 3.5 months of age. The mice were housed on a 12-hour light/dark cycle and had free access to food and water. All procedures were conducted in accordance with guidelines established by the Canadian Council on Animal Care and protocols approved by the Sunnybrook Research Institute Animal Care Committee.

#### 2.2. Experimental design

To assess the effects of FUS-BBB modulation on the proliferation of OPCs, mice (n=36) received a unilateral treatment to the left hippocampus. Mice were sacrificed at 1, 4, 7, 10, and 30 days (D) post-FUS. Mice sacrificed at 30D received bromodeoxyuridine (BrdU) (50 mg/kg, i.p.) twice daily, 2 hours apart, for the first four days post-FUS (Supplementary Fig. 1A). An independent experimental paradigm assessed the influence of FUS-BBB modulation on oligodendrogenesis in a second brain region, the striatum. Mice (n=12) received a unilateral FUS treatment to the left striatum, and animals were sacrificed at 7D or 30D post-FUS. Mice received 5-ethynyl-2’-deoxyuridine (EdU; 50 mg/kg, i.p.) once daily for the first seven days post-FUS to label cell proliferation (Supplementary Fig. 1B). EdU and BrdU have been shown to label the same populations of proliferating cells [33].

#### 2.3. Magnetic resonance imaging guided focused ultrasound

In anesthetized mice (5% isoflurane initially, reduced to 2%), depilatory cream was used to remove fur from the head, and a 26-gauge angiocatheter was inserted into the tail vein. Mouse brains were mapped using T2-weighted (T2w) MRI axial scans (7.0-T MRI [Bruker]) and four 1-mm hippocampal focal spots, or two 1-mm striatal focal spots were selected for FUS-BBB permeabilization. Coinciding with FUS sonication, mice received a tail vein injection of Definity microbubbles (0.02 ml/kg; Lantheus Medical Imaging) and Gadovist (0.2 ml/kg, Schering AG). FUS treatment was conducted using a pre-clinical MRI-guided system (an in-house developed prototype of LP100, FUS Instrument Inc., Toronto, ON, Canada), which utilizes an in-house manufactured, spherically focused transducer (1.68 MHz for hippocampal targeting or 1.78 MHz for striatal targeting, 75 mm diameter, 60 mm radius of curvature), with standard BBB opening parameters (10 ms bursts, 1 Hz burst repetition frequency, 120s sonication duration). Acoustic pressure was incrementally increased while scattered signals from the microbubbles were monitored with a polyvinylidene fluoride hydrophone [34]. When subharmonic frequencies were detected, the pressure was reduced to 25% in the hippocampus, or 50% in the striatum, where it was maintained throughout the remainder of the sonication [34]. BBB permeability was confirmed via Gadovist enhancement on follow-up T1 weighted (T1w) MRI scans. Enhancement did not differ between brain regions targeted (Supplementary Fig. 2). MRI images were analyzed using MIPAV software (v10).

#### 2.4. Tissue collection

At experimental endpoints, mice were deeply anesthetized with ketamine (150mg/kg) and xylazine (10mg/kg) and were transcardially perfused with ice-cold saline followed by 4% paraformaldehyde (PFA) in 0.1 M phosphate buffer. Whole brains were removed and post-fixed in 4% PFA for 24 hours, then transferred to 30% sucrose in 0.1 M phosphate buffer for a maximum of 72 hours. Whole brains were mounted to a sliding microtome and sectioned into 40 um-thick sections. Sections were stored in cryoprotective glycerol in multi-well plates at -20°C.

#### 2.5. Immunofluorescence staining

Four serial hippocampal sections (1:24) and six serial striatal sections (1:8) were analyzed. Sections were stained using standard immunofluorescence staining protocols. Phosphate-buffered saline (1X) was used for rinsing, antigen retrieval was performed using 10 mM sodium citrate at 80°C for 30 minutes or Tris-EDTA buffer at 70°C for 1 hour, and 5% donkey serum with 0.3% Triton-X were used for blocking. For BrdU staining, sections were antigen retrieved in 2 N HCl at 37°C for 30 minutes, rinsed, and rehydrated with 0.1 M Borate Buffer (pH=10.0) for 10 minutes. EdU staining was performed using the Click-iT EdU Cell Proliferation Kit for Imaging (Thermofisher) as per the manufacturer’s instructions. Sections were incubated with primary antibodies for 72 hours at 4°C. Primary antibodies included goat anti-GFAP (Santacruz Biotech, cat: sc-6170; 1:250), rat anti-Ki67 (ThermoFisher, cat: 14-5698-82, 1:400), rabbit anti-SOX2 (Abcam, cat: ab97959; 1:500), goat anti-OLIG2 (R&D Systems, cat: AF2418; 1:500), rabbit anti-TPPP (Abcam, cat: ab92305, 1:400), and rat anti-BrdU (Abcam, cat: ab6326; 1:400). Thereafter, sections were rinsed and incubated in secondary antibodies for 1 hour at room temperature (Jackson Immunoresearch; 1:200) or overnight at 4°C (1:400). Sections were subsequently rinsed and mounted on slides in polyvinyl alcohol-DABCO.

#### 2.6. Counting

For the first experimental paradigm, cells were counted at 63x oil objective (NA=1.4) on a Zeiss AxioImager M2 microscope. Proliferating OPCs (OLIG2^+^/Ki67^+^) and non-oligodendrocyte lineage cells (OLIG2^-^/Ki67^+^) were counted in the hippocampus between 1D and 10D post-FUS. At 30D post-FUS, mature oligodendrocytes (TPPP^+^/OLIG2^+^/BrdU^+^), immature oligodendrocytes (TPPP^-^/OLIG2^+^/BrdU^+^) and non-oligodendrocyte lineage cells (TPPP^-^/OLIG2^-^ /BrdU+) were counted. The granular and subgranular cell layers were excluded when counting proliferating non-oligodendrocyte lineage cells between 1 and 10D post-FUS and newborn non-oligodendrocyte cells at 30D to avoid neuronal-progenitor cells in this region, which are known to proliferate post-FUS. Total cell counts were calculated from four slices per animal, taken a distance of 960 um apart (1:24 sampling), and spanning a total of 2920um within the hippocampus; the average was multiplied by 72, which represents the number of 40-um slices between and including the first and last sampled slice. Only cells with a nucleus greater than 7um, corresponding the average size of the oligodendrocyte nucleus [35], were counted.

For the second experimental paradigm, single plane 20x coronal images of the striatum were obtained using a Zeiss Z1 Observer/Yokogawa spinning disk (Carl Zeiss) microscope. OPCs (OLIG2^+^/EdU^+^), immature oligodendrocytes (TPPP^-^/OLIG2^+^/EdU^+^), mature oligodendrocytes (TPPP^+^/OLIG2^+^/EdU^+^) and non-oligodendrocyte lineage cells (TPPP^-^/OLIG2^-^/EdU^+^) were quantified using a custom pipeline (Supplementary Table 1) in Cell Profiler 3.1.9. This pipeline was validated against manual counts by an experimenter blind to the experimental conditions using one image per mouse for a total of 12 images. Out of the total number of cells that were analyzed (both positive and negative), 0.69% of EdU^+^ cells, 0.35% of OLIG2^+^/EdU^+^ cells, 0.48% TPPP^+^/OLIG2^+^/EdU^+^ cells were falsely determined as positive cells for their respective staining. Total cell counts were calculated from six slices per animal, taken a distance of 320 um apart (1:8 sampling), and spanning a total of 1640um within the striatum; the average was multiplied by 40, which represents the number of 40-um slices between and including the first and last sampled slice. Only cells with a nucleus greater than 7um, corresponding the average size of the oligodendrocyte nucleus [35], were counted.

### 2.7. Statistical analysis

GraphPad Prism was used for statistical analysis. To satisfy the normality requirement for parametric testing, for the time-course analysis of proliferating OPCs and non-oligodendrocyte lineage cells (Fig. 1), raw data of cell counts was log10 transformed for statistical analysis. Subsequently, a two-way ANOVA followed by a Tukey post-hoc analysis on log10-transformed data was used to model the time-dependent and treatment-associated changes in OPC proliferation. To evaluate subject-specific increases in OPC proliferation post-FUS, a one-tailed, paired student’s t-test was performed on raw cell counts from proliferating hippocampal OPCs at 1D and 4D post-FUS (Supplementary Fig. 3). For remaining data comparing FUS-treated and untreated oligodendrogenesis at 30D post-FUS in the hippocampus, and FUS-treated and untreated OPC proliferation and oligodendrogenesis in the striatum, normality testing was conducted to assess the distribution of data using the Shapiro-Wilk test. For data that followed a normal distribution, a parametric paired Student’s t-test (one-tailed) was used to evaluate differences between conditions. If the data did not follow a normal distribution, a non-parametric paired one-tailed t-test with Welch’s correction was performed to account for unequal variances. Raw data of cell counts is plotted in Fig. 1 to Fig. 4.

**Fig 1.**
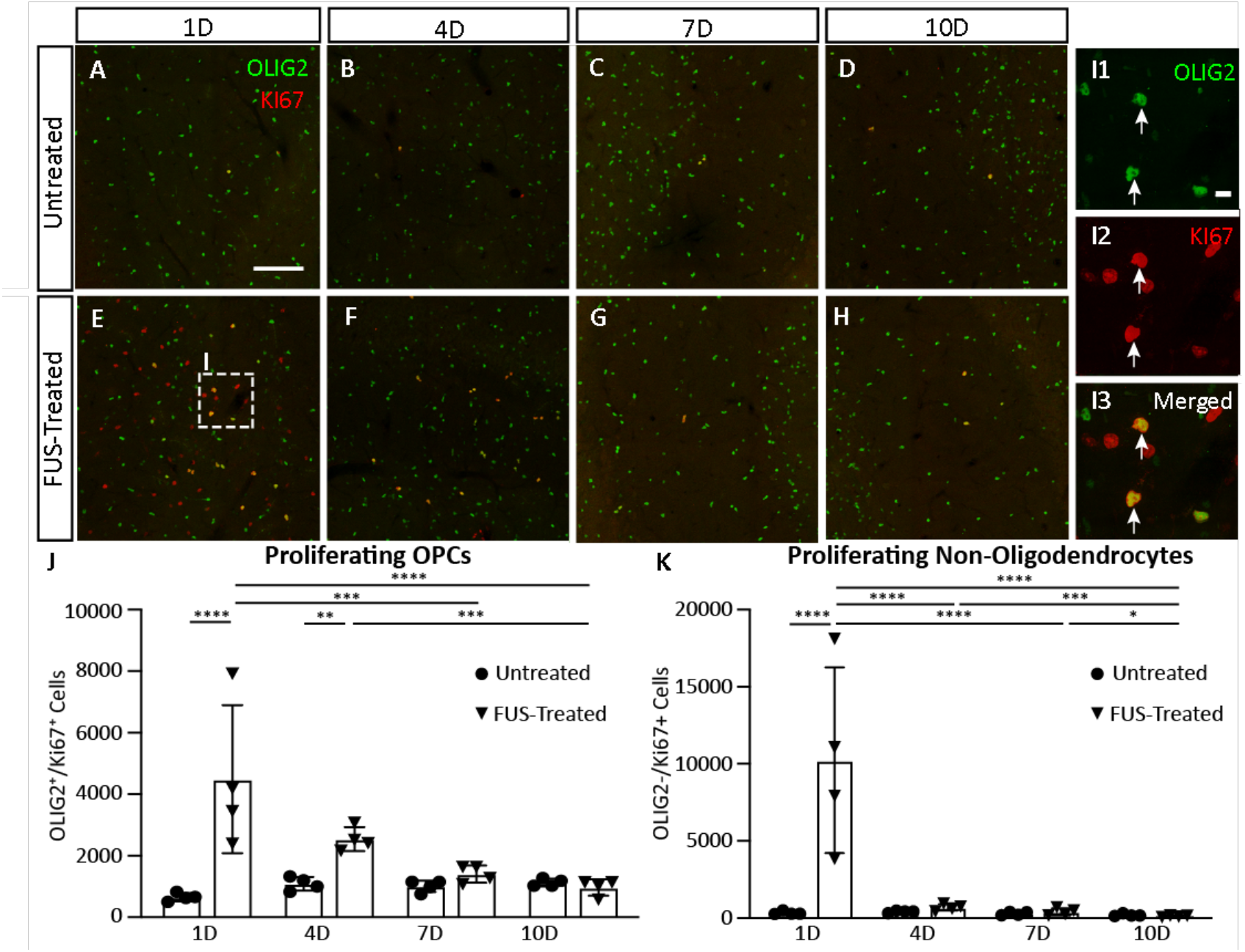
FUS-BBB modulation stimulates the proliferation of OPCs in the hippocampus. (**A-I**) Representative images of brain sections labelled for proliferating OPCs (OLIG2^+^/KI67^+^) and non-oligodendrocyte lineage cells (OLIG2^-^/KI67^+^) within (**A-D**) untreated and (**E-I**) treated hippocampi at 1, 4, 7 and 10D post-FUS. (**Insets I; I1-3**) Proliferating OPCs (arrows) and non-oligodendrocyte cells at higher magnification. (**J**) Proliferating OPCs and (**K**) non-oligodendrocytes were quantified in the untreated (circles) and FUS-treated (triangles) hippocampi at 1, 4, 7, and 10D post-FUS. Each data point is an estimate of the total cell number per unilateral hippocampus for each animal based on cell counts in four serial hippocampal sections (1:24). Statistical significance was evaluated with a two-way ANOVA followed by a Tukey post-hoc analysis on log10-transformed data. *p<0.05, **p<0.01, ***p<0.001. ****p<0.0001. Scale bars: A-D, F-I (20x) 100 um; E1-3 (63x) 10 um. D, day; FUS, focused ultrasound; OLIG2, oligodendrocyte transcription factor 2; OPC, oligodendrocyte progenitor cell.

## 3. Results

### 3.1. FUS-BBB modulation promotes hippocampal oligodendrogenesis

To assess the potential of FUS-induced oligodendrogenesis, mice received a single application of FUS, unilaterally targeting the hippocampus. Animals were sacrificed at 1, 4, 7 or 10D post-FUS and brain sections were stained with the oligodendrocyte-lineage marker, oligodendrocyte transcription factor 2 (OLIG2, green) [27] and the proliferative marker Ki67 (red) (Fig. 1A-I). Proliferating OPCs (OLIG2^+^/Ki67^+^) were quantified in the at 1, 4, 7, and 10D post-FUS (Fig. 1J), comparing the treated (triangles) and untreated (circles) hippocampi.

The average number of proliferating OPCs was 6.8-fold higher in the FUS-treated (4491±2402; mean±SD) compared to the untreated (657±133) hippocampi at 1D (Fig. 1J, ****p<0.0001) and 2.3-fold at 4D in the FUS-treated (2543±388) compared to the untreated (1094±222) side (Fig. 1J, **p=0.0027). A remarkable finding is that in all animals, the number of proliferating OPCs is increased in the FUS-treated hippocampus relative to its contralateral, untreated side at 1D and 4D post-FUS (Supplementary Fig. 3). No significant differences were detected in the number of proliferating OPCs between FUS-treated and untreated hippocampi at 7D and 10D. A significant interaction was found between time post-FUS and FUS-treatment on the proliferation of OPCs (F_(3, 24)_=8.08, ***p=0.0007).

The number of proliferating non-oligodendrocyte cells (OLIG2^-^/Ki67^+^) was 32-fold higher in the FUS-treated hippocampi (10233±6028) compared to the untreated side (320±111) at 1D (Fig. 1K, ****p<0.0001). No significant differences were observed in the number of non-oligodendrocytes between FUS-treated and untreated contralateral hippocampi at 4D, 7D, and 10D.

Next, we asked whether proliferating OPCs post-FUS differentiated into mature oligodendrocytes, which are known to express tubulin polymerization promoting protein (TPPP) [36]. To label dividing cells, animals were injected with bromodeoxyurdine (BrdU) starting at 1D and ending at 4D post-FUS. BrdU^+^ (red) cells in sections also stained with TPPP (green) and OLIG2 (blue) were counted at 30D in untreated (Fig. 2A, B) and FUS-treated (Fig. 2C-E) hippocampi 30D after FUS-BBB modulation, the number of immature oligodendrocytes (Fig. 2F) was 4.7-fold higher in the treated (5472, mean) compared to untreated (1161) hippocampi (***p=0.0004). Mature oligodendrocytes (Fig. 2G) increased by 5.3-fold in the treated (882) compared to untreated (167), hippocampi (**p=0.0027). The number of non-oligodendrocytic cells (Fig. 2H) was 5.7-fold higher in the treated (20624) compared to untreated (3632) hippocampi (***p=0.0006). These data demonstrate that FUS significantly promotes cell proliferation, including for OPCs, and leads mature oligodendrocytes to complete the process of oligodendrogenesis.

**Fig 2.**
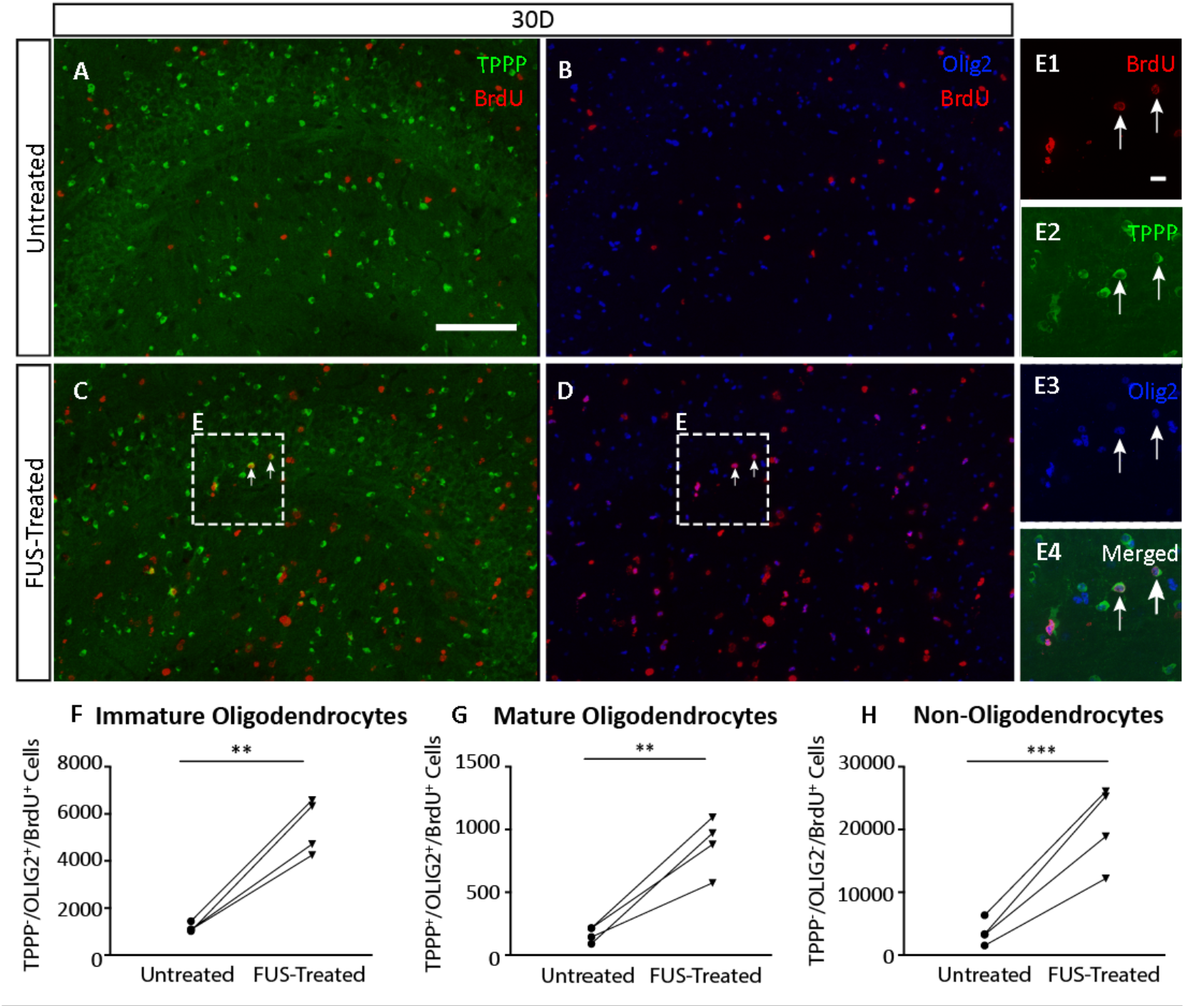
FUS-BBB modulation promotes hippocampal oligodendrogenesis. (**A-D**) Representative images of hippocampal sections labelled with antibodies against TPPP, OLIG2 and BrdU to label oligodendrocyte lineage cells in the (**A-B**) untreated and (**C-D**) FUS-treated hemispheres at 30D post-FUS. (**Insets E; E1-4**) Higher magnification showing mature oligodendrocytes (TPPP^+^/OLIG2^+^/BrdU^+^, arrows). The number of (**F**) immature oligodendrocytes (TPPP^-^/OLIG2^+^/BrdU^+^), (**G**) mature oligodendrocytes (TPPP^+^/OLIG2^+^/BrdU^+^), and (**H**) non-oligodendrocytes (TPPP^-^/OLIG2^-^/BrdU^+^) were quantified in the untreated (circles) and FUS-treated (triangles) hippocampi at 30D post-FUS Each data point is an estimate of the total cell number per unilateral hippocampus for each animal based on cell counts in four serial hippocampal sections (1:24). Statistical significance was evaluated with a one-tailed, paired student T-test. **p<0.01. ***p<0.001. Scale bars: A-D (20x) 100um; E (63x) 10um. BrdU, bromodeoxyurdine; D, day; FUS, focused ultrasound; OLIG2, oligodendrocyte transcription factor 2; TPPP, tubulin polymerization promoting protein.

### 3.2. FUS-BBB modulation stimulates OPC proliferation and oligodendrogenesis in the striatum

A different team member carried out a set of independent experiments to establish the reproducibility of whether FUS-BBB modulation induces oligodendrogenesis in another brain region. The striatum was targeted unilaterally, and different methods were used to label and quantify proliferative cells. The marker 5-ethynyl-2’-deoxyuridine (EdU) was injected intraperitoneally, once daily, from 1D to 7D a period of robust proliferation post-FUS [37,38], Animals were sacrificed 2 hours following the last EdU injection on day 7.

The colocalization of OLIG2 (green) and EdU (red) was used to label OPCs in brain sections containing the FUS-treated (Fig. 3A-D) and untreated contralateral (not shown) striatum. 7D after FUS-BBB modulation, the number of cells that proliferated and incorporated EdU (Fig.3E; EdU^+^) was 9.2-fold higher in the FUS-treated (8527) compared to the untreated (926) striatum (*p=0.0156). A 3.9-fold increase in the number post-FUS proliferating OPCs (Fig. 2F; OLIG2^+^/EdU^+^) was found in the FUS-treated (2437) compared to untreated (631) striatum (**p=0.0048). Cells that were classified as non-oligodendrocytes (Fig. 3G, OLIG2^-^/EdU^+^) were 21-fold higher in the FUS-treated (6090) compared to untreated (295) striatum (*p=0.0156). Thus, FUS-BBB modulation induces a robust increase in cell proliferation in the striatum, which includes a significant number of proliferating OPCs.

**Fig 3.**
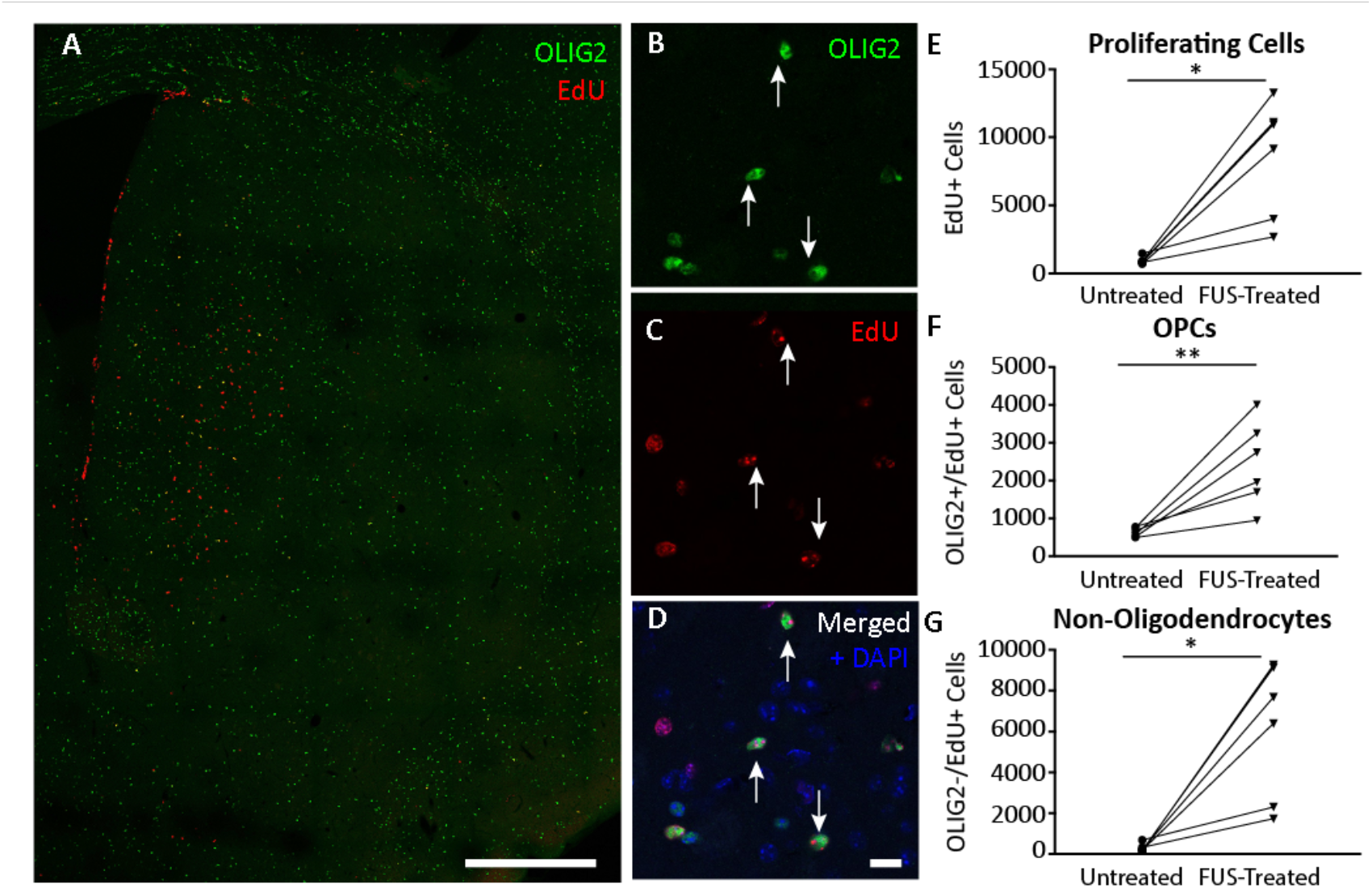
FUS-BBB modulation enhances the proliferation of OPCs in the striatum. (**A-D**) At 7D post-FUS, brain sections were stained for OPCs that had proliferated and incorporated EdU (OLIG2^+/^EdU^+^). Counts were done at higher magnification for colocalization of (**B**) OLIG2^+^ (**C**) EdU^+^, and (**D**) DAPI^+^ cells (**B-D**, white arrows). a merged image. The number of proliferating (**E**) cells (EdU^+^), (**F**) OPCs (OLIG2^+^/EdU^+^) and (**G**) non-oligodendrocytes (OLIG2^-^/EdU^+^) were quantified in the untreated contralateral (circles) and FUS-treated (triangles) striatum at 7D post-FUS. Each data point is an estimate of the total cell number per striatum as estimated by sampling six serial (1:8) striatal sections per animal. Statistical significance was evaluated with a one-tailed, paired student T-test. *p<0.05. **p<0.01. Scale bars: A (20x) 500um; B-D (63x) 10um. DAPI, 4′,6-diamidino-2-phenylindole; D, day; EdU, 5-Ethynyl-2-deoxyuridine; FUS, focused ultrasound; OLIG2, oligodendrocyte transcription factor 2; OPC, oligodendrocyte progenitor cell.

To evaluate further the process of striatal oligodendrogenesis, animals were injected with EdU from 1 to 7D post-FUS and sacrificed after 30D. Co-staining for mature oligodendrocytes (TPPP, green), immature oligodendrocytes (OLIG2, blue), and proliferating cells (EdU, red) was done at 30D post-FUS between the untreated and FUS-treated striatum (Fig. 4A-E). A trend towards an increase in the number of immature oligodendrocytes (TPPP^-^/OLIG2^+^/EdU^+^) in the FUS treated (676) compared to untreated (452) striatum was noted (Fig. 4F; p=0.0606), Dividing cells in the striatum led to a significant 5.2-fold increase in the number of mature oligodendrocytes (TPPP^+^/OLIG2^+^/EdU^+^) in the FUS-treated (1179) compared to the untreated (225) striatum (Fig. 4G;*p=0.0104). A 7.5-fold increase in the number of non-oligodendrocytes (TPPP^-^/OLIG2^-^/EdU^+^) was measured in the FUS-treated (3387) compared to the untreated (452) striatum (Fig. 4H; **p=0.0077). These data demonstrate that FUS-BBB modulation significantly promoted cell proliferation and oligodendrogenesis in the striatum.

**Fig 4.**
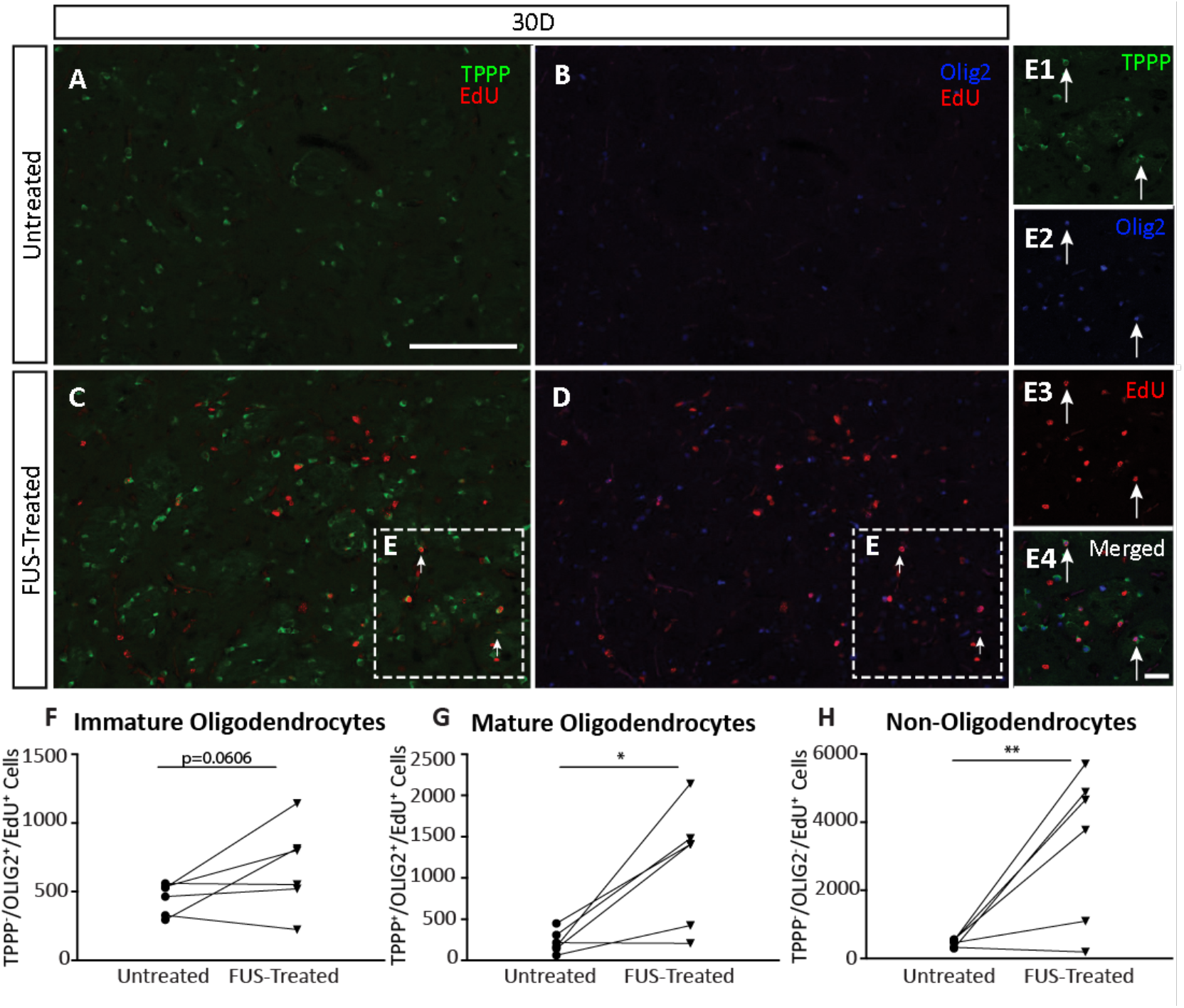
FUS-BBB modulation promotes striatal oligodendrogenesis. (**A-D**) Representative images of striatal sections stained with antibodies against TPPP, OLIG2 and EdU to label oligodendrocytes in the (**A-B**) untreated and (**C-D**) FUS-treated striatum at 30D post-FUS. (**Insets E; E1-4**) Higher magnification showing mature oligodendrocytes (TPPP^+^/OLIG2^+^/EdU^+^, arrows). The number of proliferating (**F**) immature oligodendrocytes (TPPP^-^/OLIG2^+^/EdU^+^), (**G**) mature oligodendrocytes (TPPP^+^/OLIG2^+^/EdU^+^) and (**H**) non-oligodendrocytes (TPPP^-^/OLIG2^-^/EdU^+^) were quantified in the untreated (circles) and FUS-treated (triangles) striatum at 30D. Each data point is an estimate of the total cell number per striatum as estimated by sampling six serial (1:8) striatal sections per animal. Statistical significance was evaluated with a one-tailed, paired student T-test. P-values: *p<0.05. **p<0.01. Scale bars: **A-D** (20x) 100um; **E** (63x) 10um. D, day; EdU, 5-Ethynyl-2-deoxyuridine; FUS, focused ultrasound; OLIG2, oligodendrocyte transcription factor 2; TPPP, tubulin polymerization promoting protein.

## 4. Discussion

FUS-BBB modulation has been shown to support neurobiological repair and regeneration, including adult hippocampal neurogenesis [19,22]. Furthermore, FUS-BBB modulation transiently activates astrocytes and microglia in rodent models [39], with beneficial effects such as reducing amyloid pathology for up to two weeks in the TgCRND8 mouse model of AD [19,20,22,38–41]. Here, we describe novel findings demonstrating that FUS-BBB modulation promotes oligodendrogenesis.

Specifically, FUS-BBB modulation induced a significant proliferation of OPCs which resulted in increases in oligodendrogenesis of 5.3-fold in the hippocampus and 6.7-fold in the striatum. We identified a window of oligodendrocytic regeneration in response to FUS-BBB modulation by evaluating the proliferation and maturation of OPCs over one month in the hippocampus. We found that OPC proliferation peaks at 1D and 4D post-FUS and that it is no longer significant at 10D. This window of post-FUS oligodendrocytic proliferation is consistent with previous reports of an acute inflammatory response induced by BBB-modulation by FUS [42]. Specifically, microglia reactivity, measured as increased cell density using antibodies against ionized calcium-binding adapter molecule 1 (Iba1), has been observed as early as 1 hour post-FUS [43] and up to 7D post-FUS in rodents [44], and is suggested to resolve after 2 weeks [39]. Acute inflammation in the brain can result in a range of effects, some of which are protective. For example, FUS-BBB modulation induces neuroblast formation within 1 week of sonication in rodent models of health and disease [19,22,23]. Together, these data suggest that FUS-BBB modulation induces a short-term, transient window wherein the brain mobilizes glial and neurogenic regenerative responses.

The stimulation of oligodendrogenesis in both the hippocampus and striatum suggests that OPC proliferation and differentiation may be a generalized response of brain cells to FUS-BBB modulation. Nevertheless, we observed regional differences in OPC maturation. In the hippocampus, 86% of oligodendrocytes that proliferated and incorporated BrdU from 1D to 4D post FUS remained immature 30D post-FUS (i.e., maturation rate of 14%). In contrast, 60% of oligodendrocytes in the striatum that proliferated and incorporated EdU from 1D to 7D post-FUS had differentiated into mature oligodendrocytes by 30D. One explanation for such regional differences in maturity may be attributed to the source of OPCs. OPCs in the striatum are derived from both resident OPCs and neural progenitor cells in the subventricular zone (SVZ) that retain continuous oligodendrogenesis capacity into adulthood and differentiate into oligodendrocytes under physiological and pathological conditions [45–47]. The adult hippocampus is more limited in its capacity for oligodendrogenesis, and is restricted to the hippocampal subregions of the CA3 and CA4 [48]. Additionally, the existence of regional subsets of oligodendrocytes, such as those occupying the gray and white matter with differing cellular dynamics may contribute to such differences in OPC maturation, with white matter OPCs, including those occupying the striatum, having a greater rate of proliferation and differentiation [49,50]. Furthermore, OPCs may react to the unique microenvironment created in each brain region [50]. Region-specific responses to FUS-BBB modulation can be investigated further in healthy mice and in disease models to understand the mechanisms involved and identify potential regenerative opportunities.

In both the hippocampus and striatum, FUS-BBB modulation also promoted the proliferation of cells identified as non-oligodendrocytes (*i.e*., lacking expression OLIG2). Our group and others have previously found that FUS-BBB modulation increases the number of hippocampal neural progenitor cells [22], endothelial cells [51], and microglia [43]. Specifically, using a familiar FUS-BBB modulation paradigm to target the hippocampus, *Silburt et al*. demonstrated microglia proliferation at 1 and 4D post-FUS [37], occurring alongside the gain in OPCs reported herein. Thus, several cell populations are known to proliferate in response to FUS-BBB modulation, and they can contribute to the increase in the number of non-oligodendrogenic cells that we observed.

The recognition of oligodendrocytes, OPCs, and myelin as contributing factors for the pathogenesis and progression of neurodegenerative diseases and as cellular targets of traumatic brain and spinal cord injury, has highlighted the need to develop therapeutic strategies aimed at enhancing endogenous oligodendrogenesis [31]. Most treatments for traumatic white matter injury aim to halt the progressive demyelination contributing to chronic disability [52]. Similarly, for the treatment of multiple sclerosis, approved therapies aim to modulate the immune response [53]. Thus, the development of strategies to promote myelination by enhancing oligodendrogenesis could fulfill a significant unmet medical needed for the treatment of white matter disease and CNS injury. Current preclinical strategies have ranged from small molecule drugs [54] to cell replacement strategies [55]. Previous chemical screening approaches have been used to identify small molecules, such as clemastine [56] or RNS60 [57] that enhance oligodendrocyte differentiation and/or myelination of existing OPCs [58,59], but does not expand the OPC population itself. Cell replacement therapies, likewise, have been shown in preclinical settings to effectively replace regions of demyelination, and promote regeneration, but come at the cost of requiring invasive delivery [55]. To contrast these approaches, we present FUS-BBB modulation as a novel strategy that augments endogenous oligodendrogenesis and therein expands both OPC and mature oligodendrocyte populations. Moreover, in promoting OPC proliferation, FUS may further support existing strategies to reprogram OPCs [60,61], or further accentuate the remyelination effect of existing compounds.

To evaluate the robustness and generalizability of the effects FUS on oligodendrogenesis, we conducted two independent experiments, each performed by a separate scientist. Different brain regions and methodologies for cell labelling and quantification (*e.g.,* BrdU vs. EdU) were used. Our study did not evaluate potential sex differences linked to FUS-BBB-mediated oligodendrogenesis or address its status in models of aging, disease states, and demyelination. These shortcomings are all important lines of research to explore in future experiments.

In conclusion, we show that FUS-BBB modulation, traditionally viewed as a delivery tool for the delivery of intravenous therapeutics to the brain, stimulates OPC proliferation and oligodendrogenesis in the hippocampus and striatum. Evidence for FUS-mediated oligodendrogenesis in two different brain regions suggests that oligodendrogenesis may be a generalized response to FUS-BBB modulation. The treatment of neurodegenerative disorders will require a multidimensional approach to reduce broad pathological characteristics, promote regeneration, and rescue brain function. To this end, we show that FUS-BBB modulation, stimulates oligodendrogenesis, a discovery that holds promise for the adding to the tremendous regenerative potential of FUS applications.

## Declarations

## Acknowledgements

This research could not have been possible without the expert support of Shawna Rideout-Gros, and Viva Chan who assisted with the MRIgFUS experiments and animal preparation, Kelly Coultes and Melissa Rahmati for assistance with animal care, and Kristina Mikloska and Kairavi Shah who performed the MRIgFUS procedures. Finally, we thank Dr. JoAnne McLaurin, Dr. Carol Schuurmans and Dr. Cindi Morshead for their expert guidance and critical appraisal of the current work.

## Ethics approval

All procedures were conducted in accordance with guidelines established by the Canadian Council on Animal Care and protocols approved by the Sunnybrook Research Institute Animal Care Committee.

## Availability of data and materials

Data and materials are available from the corresponding author on reasonable request.

## Authors’ contributions

KN and JS helped to conceive of the study and conducted all the analysis included in the study. KN prepared all figures. IA helped to conceive of the study and provided oversight for all analysis conducted. KH helped to conceive of the study and provided the expertise and technical support for all FUS experiments. KC and MT and provided support for animal care and collecting and processing tissues. KN, JS and IA wrote the manuscript. All authors contributed, read, provided critical analysis for, and approved the final manuscript.

## References

[1] Hynynen K, McDannold N, Vykhodtseva N, Jolesz FA. Noninvasive MR imaging-guided focal opening of the blood-brain barrier in rabbits. Radiology 2001;220:640–6. 10.1148/radiol.2202001804.

[2] Idbaih A, Canney M, Belin L, Desseaux C, Vignot A, Bouchoux G, et al. Safety and Feasibility of Repeated and Transient Blood–Brain Barrier Disruption by Pulsed Ultrasound in Patients with Recurrent Glioblastoma. Clin Cancer Res 2019;25:3793–801. 10.1158/1078-0432.CCR-18-3643.

[3] Park SH, Kim MJ, Jung HH, Chang WS, Choi HS, Rachmilevitch I, et al. Safety and feasibility of multiple blood-brain barrier disruptions for the treatment of glioblastoma in patients undergoing standard adjuvant chemotherapy. J Neurosurg 2021;134:475–83. 10.3171/2019.10.JNS192206.

[4] Wu S-K, Tsai C-L, Huang Y, Hynynen K. Focused Ultrasound and Microbubbles-Mediated Drug Delivery to Brain Tumor. Pharmaceutics 2020;13:15. 10.3390/pharmaceutics13010015.

[5] Mainprize T, Lipsman N, Huang Y, Meng Y, Bethune A, Ironside S, et al. Blood-Brain Barrier Opening in Primary Brain Tumors with Non-invasive MR-Guided Focused Ultrasound: A Clinical Safety and Feasibility Study. Sci Rep 2019;9:321. 10.1038/s41598-018-36340-0.

[6] Rezai AR, D’Haese P-F, Finomore V, Carpenter J, Ranjan M, Wilhelmsen K, et al. Ultrasound Blood–Brain Barrier Opening and Aducanumab in Alzheimer’s Disease. N Engl J Med 2024;390:55–62. 10.1056/NEJMoa2308719.

[7] Lipsman N, Meng Y, Bethune AJ, Huang Y, Lam B, Masellis M, et al. Blood–brain barrier opening in Alzheimer’s disease using MR-guided focused ultrasound. Nat Commun 2018;9:2336. 10.1038/s41467-018-04529-6.

[8] Meng Y, MacIntosh BJ, Shirzadi Z, Kiss A, Bethune A, Heyn C, et al. Resting state functional connectivity changes after MR-guided focused ultrasound mediated blood-brain barrier opening in patients with Alzheimer’s disease. NeuroImage 2019;200:275–80. 10.1016/j.neuroimage.2019.06.060.

[9] Meng Y, Goubran M, Rabin JS, McSweeney M, Ottoy J, Pople CB, et al. Blood–brain barrier opening of the default mode network in Alzheimer’s disease with magnetic resonance-guided focused ultrasound. Brain 2023;146:865–72. 10.1093/brain/awac459.

[10] Abrahao A, Meng Y, Llinas M, Huang Y, Hamani C, Mainprize T, et al. First-in-human trial of blood–brain barrier opening in amyotrophic lateral sclerosis using MR-guided focused ultrasound. Nat Commun 2019;10:1–9. 10.1038/s41467-019-12426-9.

[11] Meng Y, Pople CB, Huang Y, Jones RM, Ottoy J, Goubran M, et al. Putaminal Recombinant Glucocerebrosidase Delivery with Magnetic Resonance – Guided Focused Ultrasound in Parkinson’s Disease: A Phase I Study. Mov Disord 2022;37:2134–9. 10.1002/mds.29190.

[12] Huang Y, Meng Y, Pople CB, Bethune A, Jones RM, Abrahao A, et al. Cavitation Feedback Control of Focused Ultrasound Blood-Brain Barrier Opening for Drug Delivery in Patients with Parkinson’s Disease. Pharmaceutics 2022;14:2607. 10.3390/pharmaceutics14122607.

[13] Kofoed RH, Aubert I. Focused ultrasound gene delivery for the treatment of neurological disorders. Trends Mol Med 2024:S147149142300285X. 10.1016/j.molmed.2023.12.006.

[14] Fisher DG, Price RJ. Recent Advances in the Use of Focused Ultrasound for Magnetic Resonance Image-Guided Therapeutic Nanoparticle Delivery to the Central Nervous System. Front Pharmacol 2019;10:1348. 10.3389/fphar.2019.01348.

[15] Meng Y, Reilly RM, Pezo RC, Trudeau M, Sahgal A, Singnurkar A, et al. MR-guided focused ultrasound enhances delivery of trastuzumab to Her2-positive brain metastases. Sci Transl Med 2021;13:eabj4011. 10.1126/scitranslmed.abj4011.

[16] Gorick CM, Breza VR, Nowak KM, Cheng VWT, Fisher DG, Debski AC, et al. Applications of focused ultrasound-mediated blood-brain barrier opening. Adv Drug Deliv Rev 2022;191:114583. 10.1016/j.addr.2022.114583.

[17] Meng Y, Kalia LV, Kalia SK, Hamani C, Huang Y, Hynynen K, et al. Current Progress in Magnetic Resonance-Guided Focused Ultrasound to Facilitate Drug Delivery across the Blood-Brain Barrier. Pharmaceutics 2024;16:719. 10.3390/pharmaceutics16060719.

[18] O’Reilly MA. Exploiting the mechanical effects of ultrasound for noninvasive therapy. Science 2024;385:eadp7206. 10.1126/science.adp7206.

[19] Burgess A, Dubey S, Yeung S, Hough O, Eterman N, Aubert I, et al. Alzheimer Disease in a Mouse Model: MR Imaging–guided Focused Ultrasound Targeted to the Hippocampus Opens the Blood-Brain Barrier and Improves Pathologic Abnormalities and Behavior. Radiology 2014;273:736–45. 10.1148/radiol.14140245.

[20] Dubey S, Heinen S, Krantic S, McLaurin J, Branch DR, Hynynen K, et al. Clinically approved IVIg delivered to the hippocampus with focused ultrasound promotes neurogenesis in a model of Alzheimer’s disease. Proc Natl Acad Sci 2020;117:32691–700. 10.1073/pnas.1908658117.

[21] Mooney SJ, Shah K, Yeung S, Burgess A, Aubert I, Hynynen K. Focused Ultrasound-Induced Neurogenesis Requires an Increase in Blood-Brain Barrier Permeability. PLOS ONE 2016;11:e0159892. 10.1371/journal.pone.0159892.

[22] Scarcelli T, Jordão JF, O’Reilly MA, Ellens N, Hynynen K, Aubert I. Stimulation of Hippocampal Neurogenesis by Transcranial Focused Ultrasound and Microbubbles in Adult Mice. Brain Stimulat 2014;7:304–7. 10.1016/j.brs.2013.12.012.

[23] Shin J, Kong C, Lee J, Choi BY, Sim J, Koh CS, et al. Focused ultrasound-induced blood-brain barrier opening improves adult hippocampal neurogenesis and cognitive function in a cholinergic degeneration dementia rat model. Alzheimers Res Ther 2019;11:110. 10.1186/s13195-019-0569-x.

[24] Hatch RJ, Leinenga G, Götz J. Scanning Ultrasound (SUS) Causes No Changes to Neuronal Excitability and Prevents Age-Related Reductions in Hippocampal CA1 Dendritic Structure in Wild-Type Mice. PLOS ONE 2016;11:e0164278. 10.1371/journal.pone.0164278.

[25] Dawson M. NG2-expressing glial progenitor cells: an abundant and widespread population of cycling cells in the adult rat CNS. Mol Cell Neurosci 2003;24:476–88. 10.1016/S1044-7431(03)00210-0.

[26] Raff MC, Miller RH, Noble M. A glial progenitor cell that develops in vitro into an astrocyte or an oligodendrocyte depending on culture medium. Nature 1983;303:390–6. 10.1038/303390a0.

[27] Valério-Gomes B, Guimarães DM, Szczupak D, Lent R. The Absolute Number of Oligodendrocytes in the Adult Mouse Brain. Front Neuroanat 2018;12:90. 10.3389/fnana.2018.00090.

[28] Wang J, Zhen Y, Yang J, Yang S, Zhu G. Recognizing Alzheimer’s disease from perspective of oligodendrocytes: Phenomena or pathogenesis? CNS Neurosci Ther 2024;30:e14688. 10.1111/cns.14688.

[29] Xiao Y, Czopka T. Myelination-independent functions of oligodendrocyte precursor cells in health and disease. Nat Neurosci 2023;26:1663–9. 10.1038/s41593-023-01423-3.

[30] Hines JH. Evolutionary Origins of the Oligodendrocyte Cell Type and Adaptive Myelination. Front Neurosci 2021;15:757360. 10.3389/fnins.2021.757360.

[31] Rossi SL, Bovenkamp DE. Are oligodendrocytes the missing link in Alzheimer’s disease and related dementia research? Mol Neurodegener 2024;19:84. 10.1186/s13024-024-00760-6.

[32] Kuhlmann T, Miron V, Cuo Q, Wegner C, Antel J, Bruck W. Differentiation block of oligodendroglial progenitor cells as a cause for remyelination failure in chronic multiple sclerosis. Brain 2008;131:1749–58. 10.1093/brain/awn096.

[33] Zeng C, Pan F, Jones LA, Lim MM, Griffin EA, Sheline YI, et al. Evaluation of 5-ethynyl-2′-deoxyuridine staining as a sensitive and reliable method for studying cell proliferation in the adult nervous system. Brain Res 2010;1319C:21–32. 10.1016/j.brainres.2009.12.092.

[34] O’Reilly MA, Hynynen K. Blood-Brain Barrier: Real-time Feedback-controlled Focused Ultrasound Disruption by Using an Acoustic Emissions–based Controller. Radiology 2012;263:96–106. 10.1148/radiol.11111417.

[35] Wolswijk G. Oligodendrocyte survival, loss and birth in lesions of chronic-stage multiple sclerosis. Brain 2000;123:105–15. 10.1093/brain/123.1.105.

[36] Lehotzky A, Lau P, Tőkési N, Muja N, Hudson LD, Ovádi J. Tubulin polymerization-promoting protein (TPPP/p25) is critical for oligodendrocyte differentiation. Glia 2010;58:157–68. 10.1002/glia.20909.

[37] Silburt J. The Glial Response to Focused Ultrasound-mediated Permeabilization of the Blood-brain Barrier: Evaluating the Potential Contribution of Microglia and Astrocytes to Pathology and Regeneration. Doctoral thesis. University of Toronto, 2021.

[38] Silburt J, Aubert I. MORPHIOUS: an unsupervised machine learning workflow to detect the activation of microglia and astrocytes. J Neuroinflammation 2022;19:24. 10.1186/s12974-021-02376-9.

[39] Jordão JF, Thévenot E, Markham-Coultes K, Scarcelli T, Weng Y-Q, Xhima K, et al. Amyloid-β plaque reduction, endogenous antibody delivery and glial activation by brain-targeted, transcranial focused ultrasound. Exp Neurol 2013;248:16–29. 10.1016/j.expneurol.2013.05.008.

[40] Xhima K, Markham-Coultes K, Nedev H, Heinen S, Saragovi HU, Hynynen K, et al. Focused ultrasound delivery of a selective TrkA agonist rescues cholinergic function in a mouse model of Alzheimer’s disease. Sci Adv 2020;6:eaax6646. 10.1126/sciadv.aax6646.

[41] Poon CT, Shah K, Lin C, Tse R, Kim KK, Mooney S, et al. Time course of focused ultrasound effects on β-amyloid plaque pathology in the TgCRND8 mouse model of Alzheimer’s disease. Sci Rep 2018;8:14061. 10.1038/s41598-018-32250-3.

[42] Todd N, Angolano C, Ferran C, Devor A, Borsook D, McDannold N. Secondary effects on brain physiology caused by focused ultrasound-mediated disruption of the blood–brain barrier. J Controlled Release 2020;324:450–9. 10.1016/j.jconrel.2020.05.040.

[43] Kovacs ZI, Kim S, Jikaria N, Qureshi F, Milo B, Lewis BK, et al. Disrupting the blood– brain barrier by focused ultrasound induces sterile inflammation. Proc Natl Acad Sci 2017;114. 10.1073/pnas.1614777114.

[44] Kovacs ZI, Burks SR, Frank JA. Focused ultrasound with microbubbles induces sterile inflammatory response proportional to the blood brain barrier opening: Attention to experimental conditions. Theranostics 2018;8:2245–8. 10.7150/thno.24181.

[45] Rafalski VA, Ho PP, Brett JO, Ucar D, Dugas JC, Pollina EA, et al. Expansion of oligodendrocyte progenitor cells following SIRT1 inactivation in the adult brain. Nat Cell Biol 2013;15:614–24. 10.1038/ncb2735.

[46] Zawadzka M, Rivers LE, Fancy SPJ, Zhao C, Tripathi R, Jamen F, et al. CNS-Resident Glial Progenitor/Stem Cells Produce Schwann Cells as well as Oligodendrocytes during Repair of CNS Demyelination. Cell Stem Cell 2010;6:578–90. 10.1016/j.stem.2010.04.002.

[47] Li Y, Chen J, Zhang CL, Wang L, Lu D, Katakowski M, et al. Gliosis and brain remodeling after treatment of stroke in rats with marrow stromal cells. Glia 2005;49:407–17. 10.1002/glia.20126.

[48] DeFlitch L, Gonzalez-Fernandez E, Crawley I, Kang SH. Age and Alzheimer’s Disease-Related Oligodendrocyte Changes in Hippocampal Subregions. Front Cell Neurosci 2022;16:847097. 10.3389/fncel.2022.847097.

[49] Baxi EG, DeBruin J, Jin J, Strasburger HJ, Smith MD, Orthmann-Murphy JL, et al. Lineage tracing reveals dynamic changes in oligodendrocyte precursor cells following cuprizone-induced demyelination. Glia 2017;65:2087–98. 10.1002/glia.23229.

[50] Sherafat A, Pfeiffer F, Nishiyama A. Shaping of Regional Differences in Oligodendrocyte Dynamics by Regional Heterogeneity of the Pericellular Microenvironment. Front Cell Neurosci 2021;15:721376. 10.3389/fncel.2021.721376.

[51] McMahon D, Mah E, Hynynen K. Angiogenic response of rat hippocampal vasculature to focused ultrasound-mediated increases in blood-brain barrier permeability. Sci Rep 2018;8:12178. 10.1038/s41598-018-30825-8.

[52] Huntemer-Silveira A, Patil N, Brickner MA, Parr AM. Strategies for Oligodendrocyte and Myelin Repair in Traumatic CNS Injury. Front Cell Neurosci 2021;14:619707. 10.3389/fncel.2020.619707.

[53] Gacem N, Nait-Oumesmar B. Oligodendrocyte Development and Regenerative Therapeutics in Multiple Sclerosis. Life 2021;11:327. 10.3390/life11040327.

[54] Deshmukh VA, Tardif V, Lyssiotis CA, Green CC, Kerman B, Kim HJ, et al. A regenerative approach to the treatment of multiple sclerosis. Nature 2013;502:327–32. 10.1038/nature12647.

[55] Thiruvalluvan A, Czepiel M, Kap YA, Mantingh-Otter I, Vainchtein I, Kuipers J, et al. Survival and Functionality of Human Induced Pluripotent Stem Cell-Derived Oligodendrocytes in a Nonhuman Primate Model for Multiple Sclerosis. Stem Cells Transl Med 2016;5:1550–61. 10.5966/sctm.2016-0024.

[56] Li Z, He Y, Fan S, Sun B. Clemastine rescues behavioral changes and enhances remyelination in the cuprizone mouse model of demyelination. Neurosci Bull 2015;31:617– 25. 10.1007/s12264-015-1555-3.

[57] Rao VTS, Khan D, Jones RG, Nakamura DS, Kennedy TE, Cui Q-L, et al. Potential Benefit of the Charge-Stabilized Nanostructure Saline RNS60 for Myelin Maintenance and Repair. Sci Rep 2016;6:30020. 10.1038/srep30020.

[58] Lariosa-Willingham KD, Rosler ES, Tung JS, Dugas JC, Collins TL, Leonoudakis D. A high throughput drug screening assay to identify compounds that promote oligodendrocyte differentiation using acutely dissociated and purified oligodendrocyte precursor cells. BMC Res Notes 2016;9:419. 10.1186/s13104-016-2220-2.

[59] Mei F, Fancy SPJ, Shen Y-AA, Niu J, Zhao C, Presley B, et al. Micropillar arrays as a high-throughput screening platform for therapeutics in multiple sclerosis. Nat Med 2014;20:954– 60. 10.1038/nm.3618.

[60] Bazarek SF, Thaqi M, King P, Mehta AR, Patel R, Briggs CA, et al. Engineered neurogenesis in naïve adult rat cortex by Ngn2-mediated neuronal reprogramming of resident oligodendrocyte progenitor cells. Front Neurosci 2023;17:1237176. 10.3389/fnins.2023.1237176.

[61] Heinrich C, Bergami M, Gascón S, Lepier A, Viganò F, Dimou L, et al. Sox2-Mediated Conversion of NG2 Glia into Induced Neurons in the Injured Adult Cerebral Cortex. Stem Cell Rep 2014;3:1000–14. 10.1016/j.stemcr.2014.10.007.

