## Supplementary Data for "The Modulation of the Blood-Brain Barrier by Focused Ultrasound Stimulates Oligodendrogenesis"

### Supplementary Information:

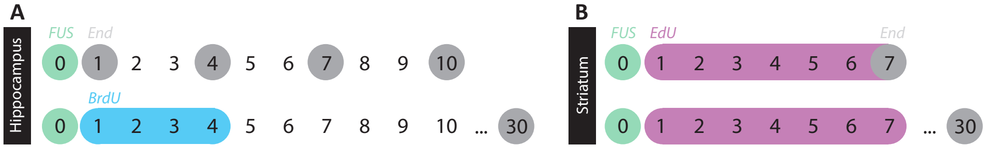


**Supplementary Fig. 1. Experimental Design.** Animals received FUS treatment targeting either the (**A**) hippocampus or (**B**) striatum. (**A**) After FUS targeting the left hippocampus on day 0, mice were sacrificed at 1, 4, 7, 10, or 30D post-FUS. Mice sacrificed at 30D received BrdU (50 mg/kg, i.p.) twice daily, 2 hours apart, for the first four days post-FUS. (**B**) Upon FUS-BBB modulation targeting the left striatum, animals were sacrificed at 7D or 30D post-FUS. Mice received EdU (50 mg/kg, i.p.) once daily for the first seven days post-FUS to label cell proliferation.
Abbreviations: BrdU; bromodeoxyuridine; EdU, 5-ethynyl-2'-deoxyuridine; FUS, focused ultrasound.

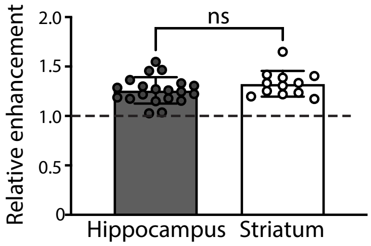


**Supplementary Fig. 2**. **Hippocampal and striatal FUS-BBB modulation.** Following FUS, T1-weighted MR images were taken to confirm increased BBB permeability at selected focal spots in the striatum or hippocampus, confirmed by Gadovist enhancement. Values greater than 1.0 indicate an increase in BBB-permeability from baseline (dotted line). Each circle presents the average relative enhancement in all focal spots per animal. There was no significant difference in enhancement level between targeting schemes. Statistical significance was evaluated with an independent student T-test.
Abbreviations: ns, not significant.

**1D post-FUS**

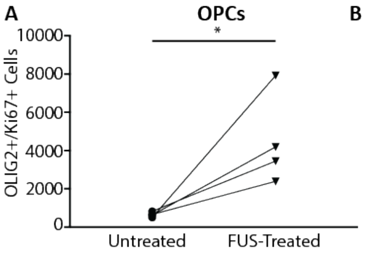


B

| Untreated | FUS-Treated |
|---|---|
| 630 | 4194 |
| 666 | 2394 |
| 828 | 3456 |
| 504 | 7920 |

**4D post-FUS**

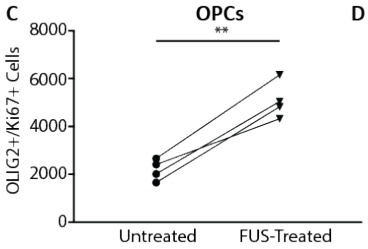


D

| Untreated | FUS-Treated |
|---|---|
| 630 | 4194 |
| 666 | 2394 |
| 828 | 3456 |
| 504 | 7920 |

**Supplementary Fig. 3. OPC proliferation is increased in the FUS-targeted hippocampus in all animals treated.** Raw data from 1 and 4D post-FUS in Fig. 1 is presented to emphasize the increase in OPC proliferation observed in each individual. At (**A, B**) 1D and (**C, D**) 4D post-FUS, all animals receiving FUS-BBB modulation targeted to the hippocampus (triangles) exhibited an increase in the number of proliferating OPCs ($OLIG2^+/Ki67^+$) compared to the contralateral untreated hippocampus (circles). Raw cell counts for each animal shown in (**B**) and (**D**) are graphed

in (**A**) and (**C**) with black lines connecting the untreated and FUS-treated hippocampus for each individual animal. Statistical significance was evaluated with a one-tailed, paired student T-test. * P<0.05, ** P<0.01.
Abbreviations: D, day; FUS, focused ultrasound; OLIG2, oligodendrocyte transcription factor 2; OPC, oligodendrocyte progenitor cell.

| Step | Module | Input | Output | Features |
|---|---|---|---|---|
| 1 | NamesAndTypes | DAPI<br>EdU<br>Olig2<br>TPPP | DAPIOrg<br>EdUOrg<br>Olig2Org<br>TPPPOrg | |
| 2 | EnhanceOrSuppressFeatures | DAPIOrg | DAPI_Enh | "Suppress" operation; Feature size 7 |
| 3 | IdentifyPrimaryObjects | DAPI_Enh | DAPI_Obj | Object diameter 6-50; adaptive two-class threshold; shape method for identifying and drawing lines between clumped objects |
| 4 | EnhanceOrSuppressFeatures | EdUOrg | EdU_Enh | "Enhance" operation; "neurite features" enhance operation; "line structures" enhancement method; feature size 50 |
| 5 | MaskImage | EdU_Enh | EdU_DAPIMsk | "DAPIObj" used as mask |
| 6 | IdentifyPrimaryObjects | EdU_DAPIMsk | EdU_DAPIMsk_Obj | Object diameter 6-60; adaptive two-class threshold; shape method for identifying and drawing lines between clumped objects |
| 7 | EnhanceOrSuppressFeatures | Olig2Org | Olig2_Enh | "Suppress" operation; Feature size 6 |
| 8 | MaskImage | Olig2Enh | Olig2_DAPIMsk | "DAPIObj" used as mask |

| | | | | |
|---|---|---|---|---|
| 9 | IdentifyPrimaryObjects | Olig2_DAPIMsk | Olig2_DAPIObj | Object diameter 6-70; adaptive two-class threshold; shape method for identifying and drawing lines between clumped objects |
| 10 | MaskImage | EdUThr | Olig2EdUDAPI_Msk | “Olig2_DAPIObj” used as mask |
| 11 | IdentifyPrimaryObjects | Olig2EdUDAPI_Msk | | Object diameter 6-70; adaptive two-class threshold; shape method for identifying and drawing lines between clumped objects |
| 12 | EnhanceOrSuppressFeatures | TPPPOrg | TPPPEnh | “Enhance” operation; “neurite features” enhance operation; “line structures” enhancement method; feature size 70 |
| 13 | Threshold | TPPPEnh | TPPPThrsh | Adaptive two-class threshold |
| 14 | MaskImage | TPPPThrsh | TPPPOlig2DAPI_Msk | “Olig2_DAPIObj” used as mask |
| 15 | EnhanceOrSuppressFeatures | TPPPOlig2DAPI_Msk | TPPPOlig2DAPI_MskEnh | “Enhance” feature type “Circles”; feature size 6 |
| 16 | IdentifyPrimaryObjects | TPPPOlig2DAPI_MskEnh | TPPPOlig2DAPI_Obj | Object diameter 6-70; adaptive two-class threshold; shape method for identifying and drawing lines between clumped objects |
| 17 | MaskImage | EdUThr | TPPPOlig2EdUDAPI_Msk | “TPPPOlig2DAPI_Obj” used as mask |
| 18 | EnhanceOrSuppressFeatures | TPPPOlig2EdUDAPI_Msk | TPPPOlig2EdUDAPIMsk_Enh | “Suppress” operation; feature size 6 |

| | | | | |
|---|---|---|---|---|
| 19 | IdentifyPrimaryObjects | TPPPOlig2EdUDAPIMsk_Enh | TPPPOlig2EdUDAPI_Obj | Object diameter 6-70; adaptive two-class threshold; shape method for identifying and drawing lines between clumped objects |
| 20 | ExportToSpreadsheet | | | |

**Supplementary Table 1. Cell Profiler pipeline for quantification of immature oligodendrocytes, mature oligodendrocytes and non-oligodendrocyte cells in the striatum.** The number of objects identified when creating EdU_DAPIMsk_Obj, Olig2EdUDAPI_Msk , and TPPPOlig2DAPI_Obj was used as the number of EdU$^{+}$, EdU$^{+}$/OLIG2$^{+,}$ and EdU$^{+}$/OLIG2$^{+}$/TPPP$^{+}$ cells, respectively.
